# Comparison of CRISPR/Cas9-mediated megabase-scale genome deletion methods in mouse embryonic stem cells

**DOI:** 10.1101/2022.03.13.484059

**Authors:** Masayuki Miyata, Junko Yoshida, Itsuki Takagishi, Kyoji Horie

## Abstract

The genome contains large functional units ranging in size from hundreds of kilobases to megabases, such as gene clusters, promoter-enhancer loops, and topologically associating domains. To analyze these large functional units, the technique of deleting the entire functional unit is effective. However, deletion of such large regions is less efficient than conventional genome editing, especially in cultured cells, and a method that can ensure success is anticipated. Here, we compared methods to delete the 2.5-Mb Krüppel-associated box zinc finger protein (KRAB-ZFP) gene cluster on chromosome 4 in mouse embryonic stem cells using CRISPR/Cas9. Three methods were used: first, deletion by non-homologous end joining (NHEJ); second, homology-directed repair (HDR) using a single-stranded oligodeoxynucleotide (ssODN) with 70-bp homology arms; and third, HDR employing targeting vectors with a selectable marker and 1-kb homology arms. NHEJ-mediated deletion was achieved in 9% of the transfected cells. The deletion frequency of NHEJ and HDR was found to be comparable when the ssODN was transfected. Deletion frequency was highest when targeting vectors were introduced, with deletions occurring in 31–63% of the drug-resistant clones. Biallelic deletion was observed when targeting vectors were used. This study will serve as a benchmark for the introduction of large deletions into the genome.

## INTRODUCTION

Recent progress in genome science has revealed large functional units in the genome— such as gene clusters, promoter-enhancer loops, and topologically associating domains—which range from several hundred kilobases to megabases in length (Merkenschlager & Nora, 2016). Dysregulation of these functional units can lead to human diseases (Hnisz *et al*, 2016). To understand the genome from the perspective of such large functional units, technologies that can reliably modify large genomic regions are required. For this purpose, megabase-scale genome modifications using CRISPR/Cas9 have been reported by either microinjection of Cas9 and gRNAs into zygotes (Boroviak *et al*, 2016; Kato *et al*, 2017; Korablev *et al*, 2017; Mizuno *et al*, 2015) or by transfection of cultured cells (Eleveld *et al*, 2021; Essletzbichler *et al*, 2014; Wolf *et al*, 2020). However, the efficiency of megabase-genome modification is inferior in cultured cells compared to zygote microinjection. Biallelic megabase-scale deletion is even more challenging in cultured cells but is unquestionably required to conduct phenotype analysis in cultured cells. Genome modification in cultured cells is particularly important in the study of human biology and diseases. This is because zygote microinjection to create genetically modified living organisms is ethically prohibited in humans; hence, cultured cells must be used for analysis. Therefore, we anticipate the establishment of an efficient protocol for megabase-scale genome modifications in cultured cells.

The distal region of mouse chromosome 4 contains a 2.5-Mb region within which a gene cluster of Krüppel-associated box zinc finger protein (KRAB-ZFP) genes resides (Wolf *et al*., 2020). KRAB-ZFP genes are known to be transcriptional repressors of retrotransposons (Ecco *et al*, 2017). The retrotransposons and KRAB-ZFP genes have diversified in both nucleotide sequence and copy number as a result of their arms race. The diversified KRAB-ZFP genes also function as regulators of endogenous genes (Ecco *et al*., 2017). We considered the deletion of this 2.5-Mb KRAB-ZFP gene cluster as an experimental model for megabase-scale genomic deletion in cultured cells and compared three methods employing the CRISPR/Cas9 system: (1) non-homologous end joining (NHEJ), (2) homology-directed repair (HDR) using an ssODN donor, and (3) HDR using double-stranded targeting vectors. The results will serve as a benchmark for megabase-scale genomic deletion methods in cultured cells.

## RESULTS

### Overview of the methods for deleting the 2.5-Mb genomic region

Figure 1 shows the 2.5-Mb genomic region of the KRAB-ZFP gene cluster located on the distal side of mouse chromosome 4. We attempted to delete this entire region in mouse embryonic stem cells (ESCs) by cleaving the upstream and downstream sites with single guide RNAs (sgRNAs).

**Figure 1.**
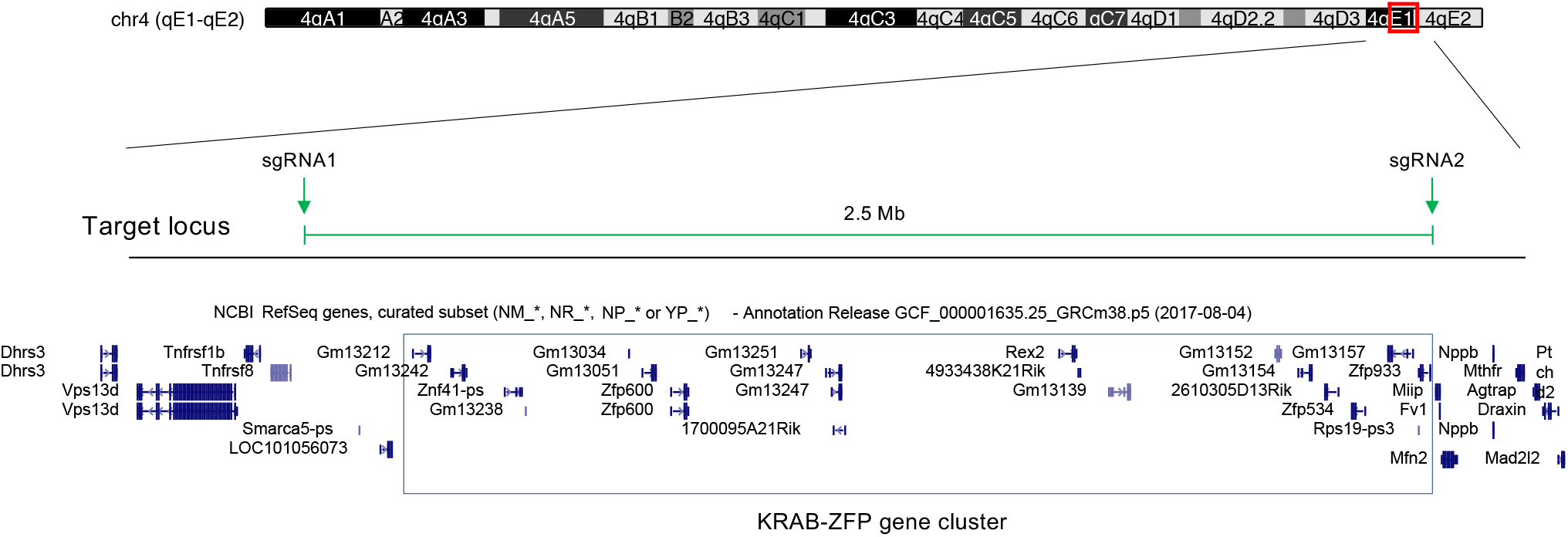
Genomic view of the KRAB-ZFP gene cluster on chromosome 4. UCSC genome browser view of the KRAB-ZFP gene cluster and the position of the sgRNAs used for genomic deletion.

We compared three methods (Fig. 2A). In all methods, the plasmid vector pX330 (Cong *et al*, 2013) was used to express Cas9 and sgRNAs, and the TransFast transfection reagent, which employs lipid-mediated gene transfer, was utilized to introduce DNA into ESCs. In Method 1, repair template DNA was not transfected. Therefore, cleaved sites were repaired by NHEJ. In Method 2, a 146-base ssODN containing 70-base 5’and 3’ homology arms (Fig. 2B) was co-transfected as a repair template for HDR. We introduced an EcoRI site between the homology arms (Fig. 2B) to facilitate the identification of HDR events. It was expected that NHEJ would still be observed in Method 2 in case the ssODN was not utilized during repair. To enrich transfected ESCs, we co-transfected a puromycin resistance gene expression vector in Methods 1 and 2 (Fig. 2A) and selected ESCs using puromycin between 24 h and 72 h after transfection (Fig. 2C, left). The ESCs were then sparsely plated on mitomycin C-treated mouse embryonic fibroblast (MEF) feeder cells. After 8 days of culture, colonies were picked and analyzed by polymerase chain reaction (PCR) (Fig. 2C, left). In Method 3, we transfected two double-stranded targeting vectors together with the Cas9/sgRNA expression vector (Fig. 2A, 2B). Each targeting vector contained the hygromycin resistance (hyg) gene and the neomycin resistance (neo) gene, respectively. Both vectors contained the same 1-kb homology arms corresponding to the upstream and downstream regions of the genomic cleavage sites. We expected that co-transfection of two targeting vectors and selection for hygromycin/G418 double-resistance would increase the efficiency of identifying biallelic deletions. One day after transfection, ESCs were split and selected with both hygromycin and G418, hygromycin only, and G418 only (Fig. 2C, right). Nine days after selection, drug-resistant colonies were picked and analyzed by PCR.

**Figure 2.**
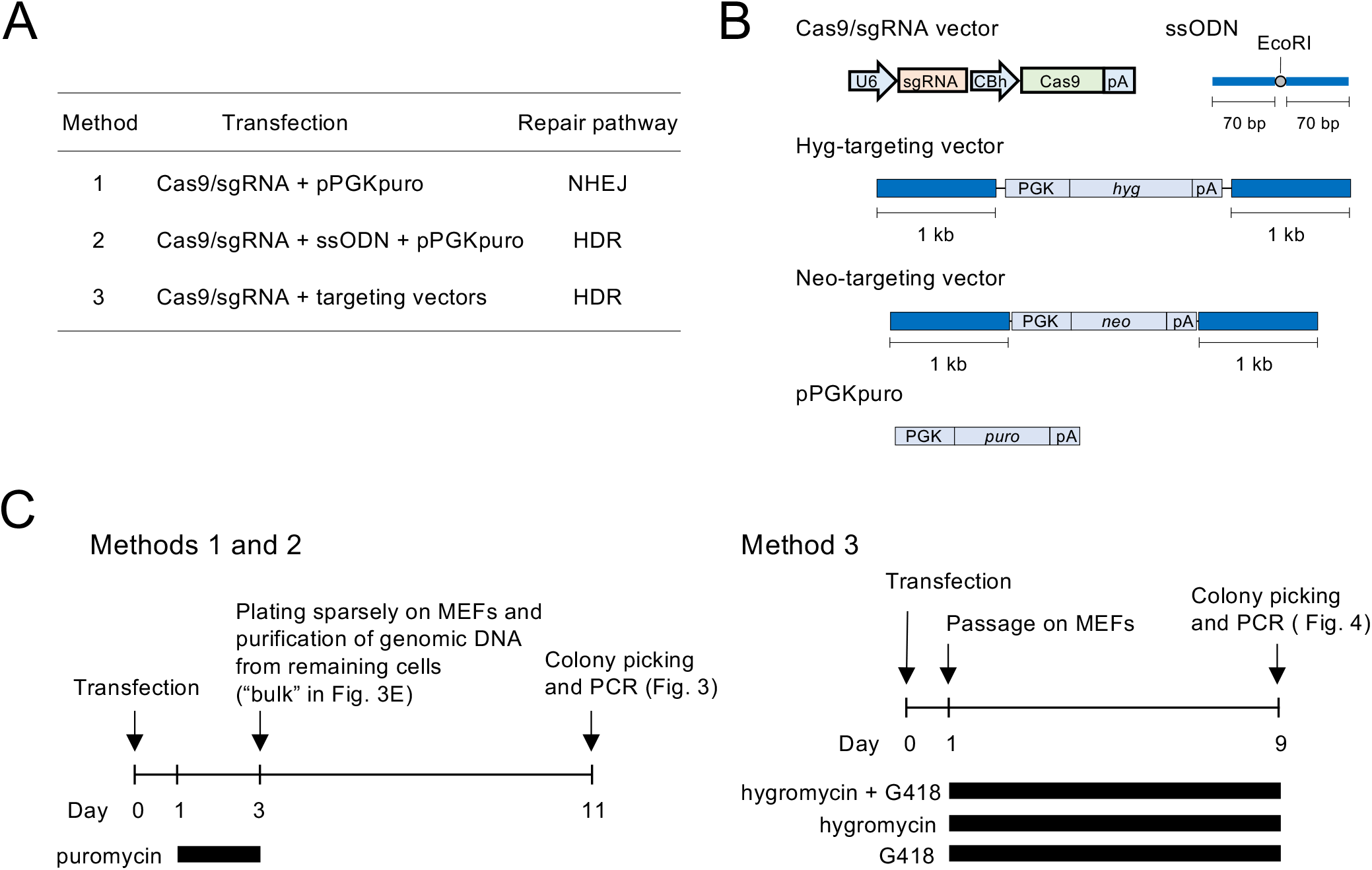
Protocols for inducing genomic deletion. (A) Summary of the three methods compared in this study. (B) Schematic of the vector structures. U6, U6 promoter; CBh, truncated CBA hybrid promoter; PGK, phosphoglycerate kinase-1 promoter; pA, polyadenylation signal. (C, D) Time course of Methods 1, 2, and 3.

### Comparison of genomic deletions with and without the ssODN

We compared Method 1, in which no repair template was introduced, and Method 2, in which an ssODN was introduced as a repair template. Single-cell–derived colonies were isolated according to the protocol in Figure 2C, and genomic deletions were detected by PCR as shown in Figure 3A. Both HDR and NHEJ were expected to occur in Method 2. We distinguished them by digesting the PCR products with EcoRI (Fig. 3A, right).

**Figure 3.**
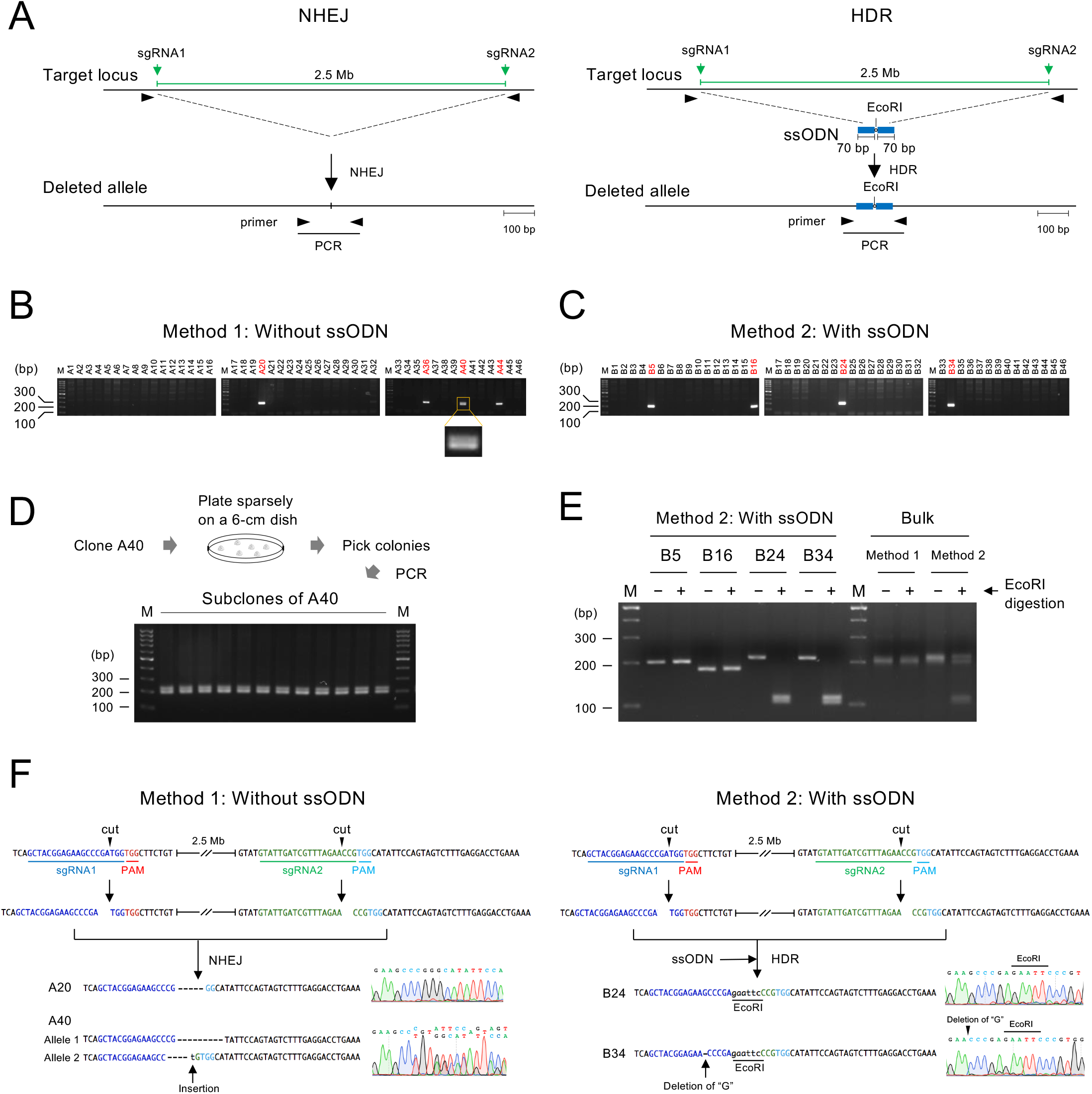
Genomic deletion induced by Method 1 and Method 2. (A) Predicted scheme of genomic deletions induced by NHEJ (left) and ssODN-mediated HDR (right). (B, C) PCR screening of genomic deletions in Method 1 (B) and Method 2 (C). A magnified view of clone A40 is shown to depict the two bands that are close in size. M, 100-bp size marker. (D) Schematic diagram showing the procedure for subcloning A40 and the result of the PCR analysis. (E) EcoRI digestion of PCR products obtained in Method 2. (F) Representative results of the sequence analysis of the PCR products obtained in Method 1 (left) and Method 2 (right). Dashed lines indicate nucleotide deletions from the Cas9/sgRNA-mediated cleavage site.

In both Method 1 and Method 2, four out of 46 colonies (9%) were PCR-positive (Fig. 3B, C). In Method 1, two bands of similar size were observed in one lane (A40 in Fig. 3B). To exclude the possibility that two clones were fused during ESC colony formation, PCR was conducted after recloning (Fig. 3D). However, the two bands were observed even after recloning. Therefore, we concluded that these two bands derive from a single clone. Their presence suggests the possibility that deletions occurred in both alleles. We address this point later in Figure 5.

To compare the efficiency of HDR and NHEJ in Method 2, the four PCR products obtained in Figure 3C were digested by EcoRI. EcoRI cleavage was observed in two of the products (Fig. 3E), indicating that the efficiency of HDR and NHEJ was comparable. To further assess this observation, we purified genomic DNA from the bulk cell population (Fig. 2C), conducted PCR to amplify deletion junctions, and digested the PCR products with EcoRI (Fig. 3E). As expected, the PCR product obtained by Method 1 was not cleaved by EcoRI. On the other hand, EcoRI-cleaved bands were observed in the products derived by Method 2, and the density of cleaved and uncleaved bands was similar. Thus, as in the analysis of cloned cells, the efficiency of HDR and NHEJ was considered comparable.

To confirm that the PCR amplification represented the deletion of the targeted gene cluster, we sequenced the PCR products (Fig. 3F). Two clones derived by both Method 1 and Method 2 were analyzed, and the results confirmed that all PCR products represented the deletion of the targeted gene cluster. Both PCR products from Method 2 were cleaved with EcoRI (Fig. 3E), suggesting that the deletion was completed in a precise manner. However, sequence analysis revealed that one of the PCR products had a single base deletion of a guanine nucleotide upstream of the EcoRI site (B34 in Fig. 3F). Since oligonucleotide synthesis is not perfectly accurate, we speculate that this deletion may have been pre-existing in the ssODN.

### Genomic deletions using targeting vectors

Next, we attempted genomic deletion by Method 3, which involves the transfection of hyg- and neo-targeting vectors for HDR. Following transfection, ESCs were selected with both hygromycin and G418, hygromycin only, or G418 only (Fig. 2C). Drug-resistant clones were analyzed by PCR using two primer pairs that detect HDR in the upstream and downstream regions (Fig. 4A).

**Figure 4.**
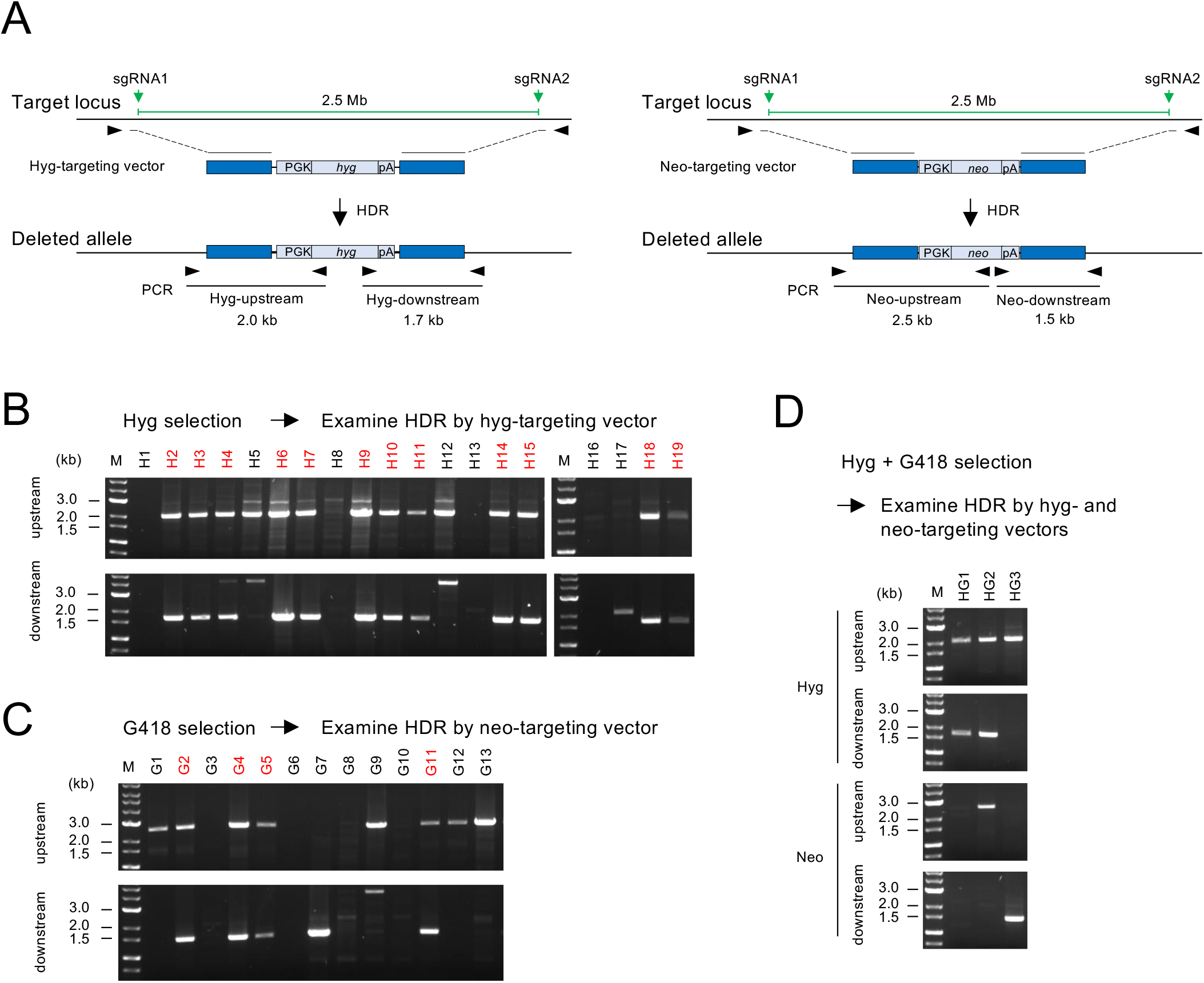
Genomic deletion induced by Method 3. (A) Predicted scheme of genomic deletions induced by HDR following transfection of targeting vectors. Two targeting vectors, each containing the hyg and the neo cassette, were co-transfected. (B-D) PCR screening of genomic deletions in hygromycin-resistant clones (B), G418-resistant clones (C), and hygromycin/G418 double-resistant clones (D). M, 1-kb size marker.

First, we analyzed hygromycin-resistant clones. Out of the 19 clones analyzed, the expected recombination was observed in both upstream and downstream regions in 12 clones (63%; Fig. 4B). Next, we analyzed 13 G418-resistant clones and observed the expected recombination in both upstream and downstream regions in 4 clones (31%; Fig. 4C). Thus, the mean deletion efficiency of HDR using targeting vectors was 47%, which is 5 times higher than that of NHEJ or ssODN-mediated HDR. Finally, we analyzed hygromycin/G418 double-resistant clones to investigate whether they harbor a biallelic mutation. We obtained much fewer colonies via hygromycin/G418 double-selection compared to the single selections. We analyzed 3 double-resistant clones by PCR using 4 primer sets for each clone; however, the expected recombination was not observed with at least one of the primer sets (Fig. 4D), suggesting that biallelic deletion is not a frequent event.

### Comparison of biallelic deletion frequency between the three methods

To analyze the phenotypes caused by genomic deletions, it is often necessary to introduce deletions in both alleles. However, the results of the analysis of hygromycin/G418 double-resistant clones suggested that biallelic deletion is not frequent (Fig. 4D). Therefore, we systematically compared the frequency of biallelic deletion among the three methods by analyzing the PCR-positive clones shown in Figures 3 and 4.

We set up two pairs of PCR primers within the deleted region (Fig. 5A). If both alleles were deleted, no amplification should be detected. For Method 1 and 2, we analyzed all clones that showed deletion in Figure 3. PCR amplification was observed in all clones (Fig. 5B), suggesting that biallelic deletion did not occur. This includes the clone A40, which showed two bands in Figures 3B and 3D. The clone A40 could be aneuploid and have, for example, three copies of chromosome 4, two having undergone deletion and one being retained intact. For Method 3, we analyzed 12 hygromycin-resistant clones in which the predicted recombination was observed in both upstream and downstream regions (Fig. 4B). No amplification was observed in three clones (H4, H11, H19; Fig. 5C), suggesting that both alleles were deleted in these clones. To investigate whether biallelic deletion accompanied NHEJ, which does not involve the recombination of the targeting vector as observed in Method 1, we conducted the same PCR analysis performed in Figure 3B. No amplification was detected (Fig. 5D), suggesting that either the NHEJ observed in Method 1 did not occur or that NHEJ with the deletion of the binding site of the PCR primer occurred. To examine the possibility that biallelic deletion involved HDR by the neo-targeting vector, we conducted the same PCR analysis shown in Figure 4C. No amplification was detected in two of the three biallelic mutants (Fig. 5E). Although PCR amplification was detected in one of the mutants in the analysis of the upstream region (clone H11), the band size was different from the expected one and no amplification was detected in the downstream region (Fig. 5E), suggesting that HDR by the neo-targeting vector did not occur. On the basis of the results of Figures 5D and 5E, we speculate that biallelic deletion was introduced through biallelic HDR by the hyg-targeting vector or through the combination of single allele HDR by the hyg-targeting vector and NHEJ accompanied by the deletion of the PCR primer binding site.

**Figure 5.**
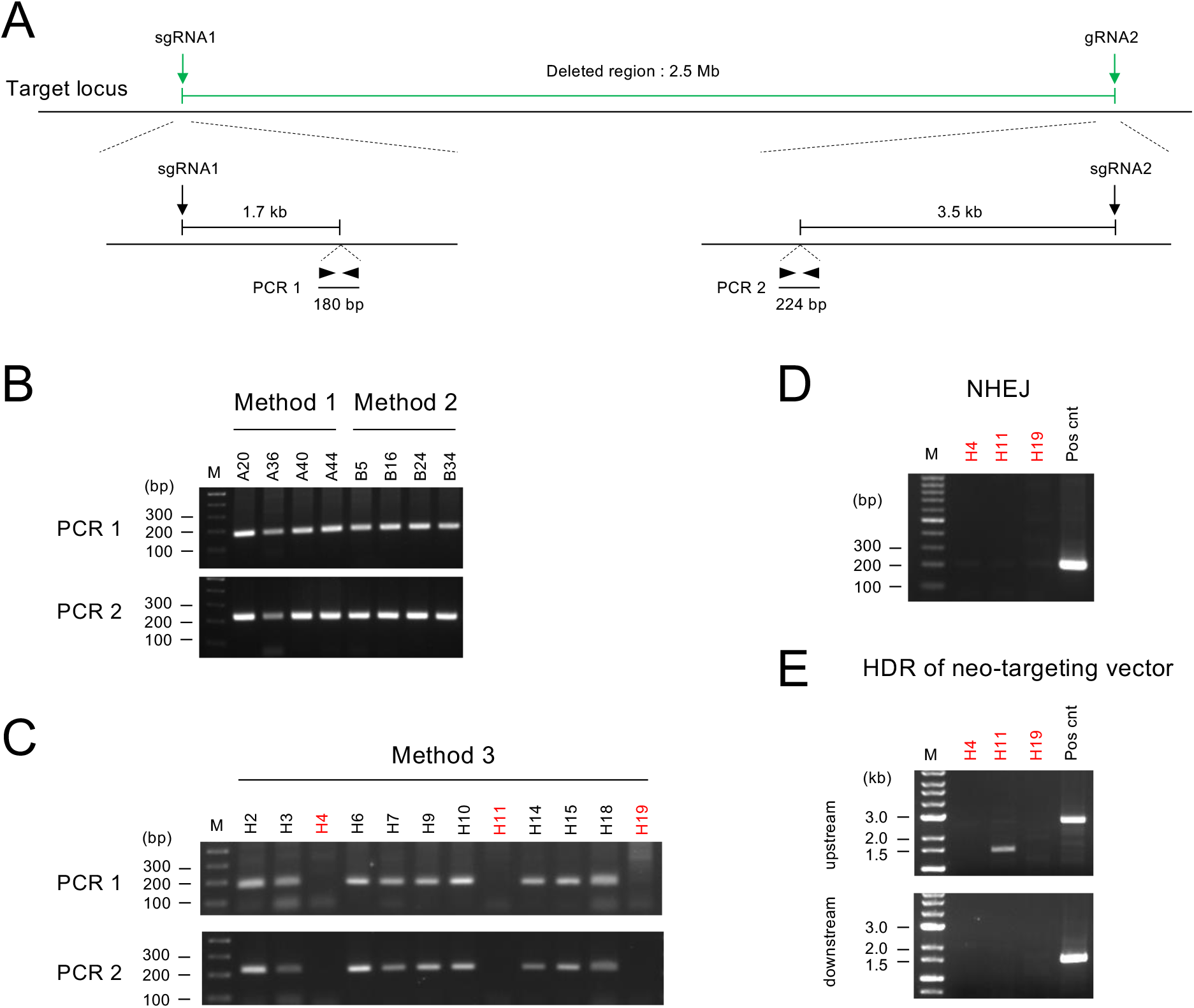
Identification of biallelic deletion events. (A) Location of the PCR primers for the screening of biallelic deletion. (B, C) PCR screening for biallelic deletion of the candidate clones obtained by Method 1 and Method 2 (B) and Method 3 (C). (D) Screening for NHEJ-mediated genomic deletion. The same PCR protocol as in Figure 3B was performed. Clone A20, which was PCR-positive in Figure 3B, was used as a positive control as indicated (Pos cnt). (E) Screening for HDR mediated by the neo-targeting vector. The same PCR protocol as in Figure 4C was performed. Clone G2, which was PCR-positive in Figure 4C, was used as a positive control (Pos cnt).

### RNA-seq analysis of deletion mutants

Although the absence of PCR amplification within the deletion target site supports biallelic deletion in the three clones H4, H11, and H19 (Fig. 5C), we could not clarify the deletion junction by PCR analysis (Fig. 5D, E). To determine whether the biallelic deletion was confined to the expected region, we performed RNA-seq in the following four cell lines—wild-type ESCs, the clone H14 with single-allele deletion, and the clones H4 and H19 with biallelic deletion— and compared the gene expression at the deletion target site (Fig. 6, Supplementary Table 1). In the single-allele deletion clone (H14), the expression of the gene cluster within the deletion target site was reduced by approximately two fold compared with wild-type ESCs, as predicted (Fig. 6A, left). By contrast, in the biallelic deletion clones (H4 and H19), the expression of the gene cluster was almost undetectable (Fig. 6A, middle and right), confirming the biallelic deletion of the 2.5-Mb gene cluster. We then analyzed the gene expression around the upstream (Fig. 6B) and the downstream (Fig. 6C) deletion junctions. The gene expression was detectable outside the deletion target site in both biallelic mutants (Fig. 6B, C), indicating that the deletion was confined to the expected region. These results demonstrate that the biallelic deletion of the 2.5-Mb gene cluster was achieved using the targeting vectors.

**Figure 6.**
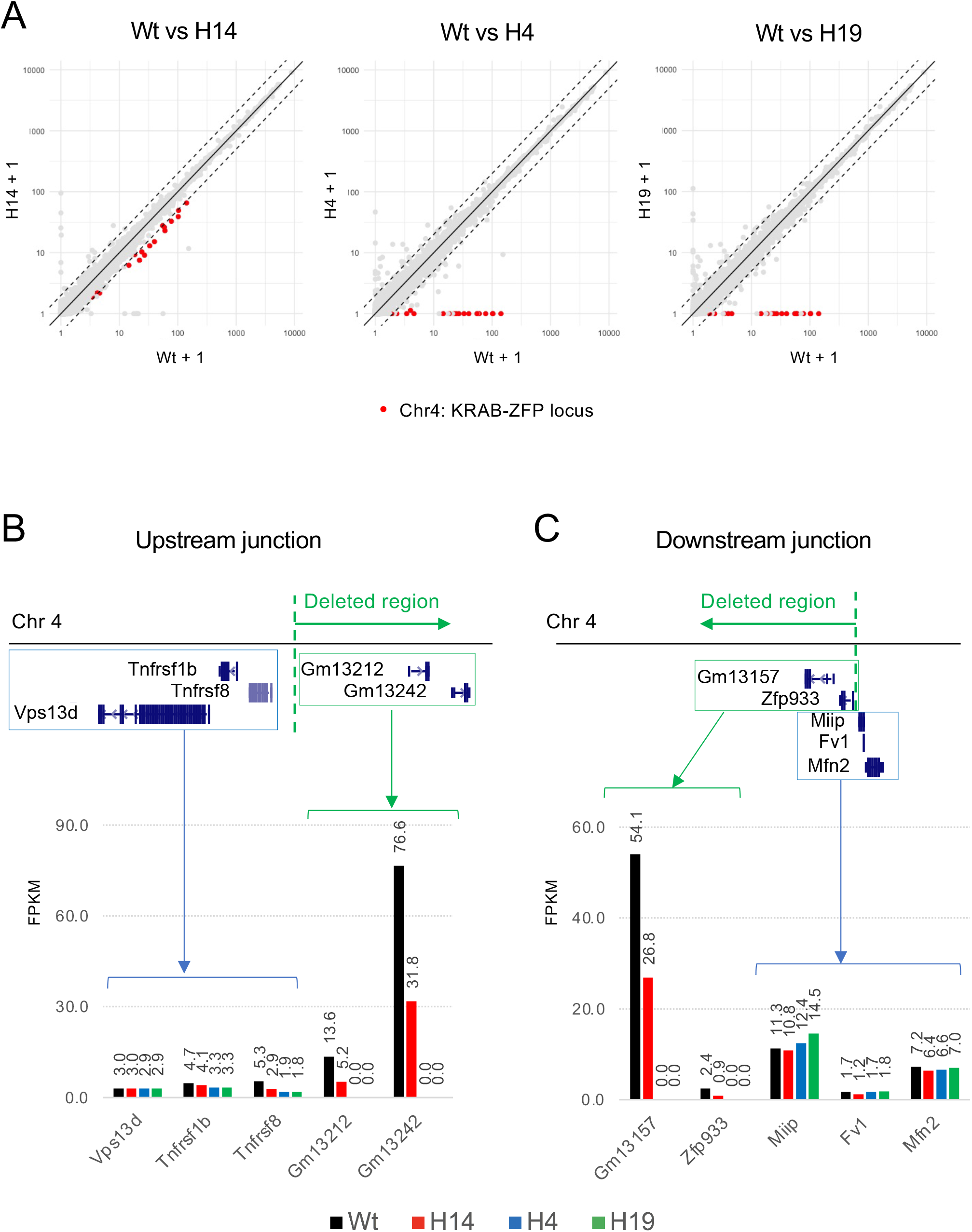
RNA-seq analysis of deletion mutants. (A) Expression analysis of the KRAB-ZFP gene cluster. The gene expressions of the single-allele deletion mutant (H14) and biallelic deletion mutants (H4 and H19) were compared with wild-type ESCs (Wt). Red dots indicate the expression of the KRAB-ZFP gene cluster under study. Data are shown in FPKM. (B, C) Gene expression at the upstream (B) and downstream (C) deletion junctions. Note that the gene expression outside the deletion target site was detectable in both biallelic deletion mutants (H4 and H19), indicating that biallelic deletion was confined to the predicted region.

## DISCUSSION

In this study, we compared three megabase-scale genomic deletion methods in mouse ESCs: Method 1 using Cas9/sgRNA only, Method 2 using Cas9/sgRNA and ssODN, and Method 3 using Cas9/sgRNA and targeting vectors. The results showed that all methods are feasible at least for monoallelic deletion. On the other hand, the three methods differed in the simplicity of the experimental design and the efficiency of deletion. Therefore, the choice of method depends on the purpose of the experiment. In the following section, we compare the three methods and discuss some considerations to further improve deletion efficiency.

The deletion efficiency in Methods 1, 2, and 3 were 9%, 9%, and 31–63%, respectively. Furthermore, biallelic deletion was observed only in Method 3. This indicates that Method 3, which uses a targeting vector, is superior to the others when considering only the efficiency of deletion. However, there are some drawbacks to using targeting vectors. First, generating a targeting vector is time-consuming. Second, setting up experimental conditions for PCR screening of the deletion clones may take time compared to Methods 1 and 2 because a longer PCR amplification is required. Therefore, Methods 1 and 2, which are straightforward in their experimental design, may be sufficiently effective if the number of clones to be screened by PCR is manageable. In fact, a recent report demonstrated biallelic deletion of the same KRAB-ZFP gene cluster by a procedure similar to Method 2 (Wolf *et al*., 2020). Although the efficiency of biallelic deletion is not described in this report, the results suggest that sufficient deletion efficiency may be achieved without using a targeting vector by optimizing experimental conditions.

Several possible improvements can be made to increase the efficiency of megabase-scale genomic deletion. The first is to use a ribonucleoprotein (RNP) complex consisting of Cas9 and sgRNA. A recent report demonstrated that transfection of a Cas9/sgRNA RNP complex was more efficient in cleaving the genome than transfection of Cas9/sgRNA expression vectors. The authors argue that the intracellular assembly of Cas9 and sgRNAs expressed from transfected vectors is hampered by the competitive binding of mRNA to Cas9 (Kagita *et al*, 2021). Second, optimization of transfection conditions may significantly affect the deletion efficiency. In the same report described above, two electroporators, MaxCyte and 4D-Nucleofector, were compared for the introduction of mutations into human iPSCs by ssODNs (Kagita *et al*., 2021). The results showed that MaxCyte was superior to 4D-Nucleofector in terms of mutagenesis efficiency. In our study, cationic lipid-based transfection reagents were used. The use of other transfection methods may improve the efficiency of megabase-scale genomic deletions. Third, the use of single-stranded targeting vectors may be useful. Previous studies in zygote microinjection have suggested that long single-stranded DNA donors are efficient templates for HDR (Codner *et al*, 2018; Miura *et al*, 2015; Quadros *et al*, 2017). However, targeting vectors used in cultured cells are usually several kb in length because of the presence of a selection marker cassette, and the preparation of such a long single-stranded DNA of high quality is labor-intensive. Recently, it has become possible to synthesize long single-stranded DNA commercially, which may apply to megabase-scale deletion.

Taken together, the results of this study will serve as a benchmark for selecting methods to introduce megabase-scale genomic deletions.

## METHODS

### Cell line and cell culture

The REC24-3 mouse ESC line, a derivative of the V6.5 mouse ESC line (Eggan *et al*, 2001), was used in this study. REC24-3 contains the ERT2-iCre-ERT2 cassette (Casanova *et al*, 2002) at the Rosa26 locus (Zambrowicz *et al*, 1997), which was introduced by the same procedure described previously (Horie *et al*, 2011). The presence of the ERT2-iCre-ERT2 cassette is irrelevant to the purpose of this study. ESCs were cultured in a serum-containing medium composed of KnockOut DMEM (Thermo Fisher Scientific) supplemented with 20% fetal bovine serum (FBS), non-essential amino acids, 0.1 mM 2-mercaptoethanol and 1,000 U/ml leukemia inhibitory factor (LIF; Millipore). Mitomycin C (MMC)-treated MEFs were used as feeder cells.

### Construction of the Cas9/sgRNA expression vectors

Cas9 and sgRNAs were expressed using pX330 (Cong *et al*., 2013). Complementary oligonucleotides for each sgRNA (Supplementary Table 2) were annealed and cloned into the BbsI site of pX330.

### Construction of the targeting vectors

The targeting vectors were constructed using the primers listed in Supplementary Table 2 as follows. A 1-kb genomic fragment upstream of the cleavage site of gRNA1 was PCR-amplified from C57BL/6J genomic DNA using the primers Zfp600-5HR1-F1 and Zfp600-5HR1-R1. The fragment was digested with KpnI and HindIII and cloned into the KpnI-HindIII site of pPGKneo-F2F (gift from Dr. K. Yusa) adjacent to the neo selection cassette, resulting in pPGKneoF2F-Zfp600-5HR. Next, a 1-kb genomic fragment downstream of the cleavage site of gRNA2 was PCR-amplified from C57BL/6J genomic DNA using primers Zfp600-3HR1-F1 and Zfp600-3HR1-R1. The fragment was digested with NotI and SacII and cloned into the NotI-SacII site of the pPGKneoF2F-Zfp600-5HR, which is located opposite to the first cloning site of the neo selection cassette, resulting in the neo-targeting vector pZfp600-DEL-TV1-Neo. The HindIII-NotI neo cassette of pZfp600-DEL-TV1 was replaced with the HindIII-NotI hyg cassette of pPGKhyg-F2F (gift from Dr. K. Yusa), resulting in the hyg-targeting vector pZfp600-DEL-TV1-Hyg.

### Transfection

The TransFast transfection reagent (Promega) was used in all transfections. ESCs (2.5 × 10^5^) were mixed with 2.5 μg of DNA and 15 μl of TransFast in serum-containing medium in a total volume of 500 μl and plated onto one well of a 24-well plate seeded with MMC-treated MEFs. After 1 h, 1 ml of medium was added to the well, and the medium was replaced with fresh medium 10 h after transfection. After this step, different culture protocols were utilized depending on the purpose of the experiment as described below.

### Comparison of the deletion protocols

#### Method 1

On Day 0, ESCs (2.5 × 10^5^) were transfected with 1.125 μg of pX330-gRNA1, 1.125 μg of pX330-gRNA2, and 0.25 μg of the puromycin resistance gene expression vector (pPGKpuro). ESCs were selected by 1 μg/ml of puromycin from Day 1 to Day 3 to enrich transfected cells. After completing puromycin selection on Day 3, ESCs were dissociated with trypsin/EDTA and plated sparsely on MEFs without puromycin for single-cell cloning; the remaining cells were subjected to genomic DNA purification as a bulk control. On Day 11, ESC colonies were picked and divided into two groups: one for PCR analysis and the other for continuous culture to make frozen stocks.

#### Method 2

On Day 0, ESCs (2.5 × 10^5^) were transfected with 0.75 μg of pX330-gRNA1, 0.75 μg of pX330-gRNA2, 0.75 μg of ssODN, and 0.25 μg of pPGKpuro. The remaining procedure is the same as in Method 1.

#### Method 3

On Day 0, ESCs (2.5 × 10^5^) were transfected with 0.625 μg of pX330-gRNA1, pX330-gRNA2, pZfp600-DEL-TV1-Hyg, and pZfp600-DEL-TV1-Neo. On Day 1, ESCs were dissociated with trypsin/EDTA, plated onto 6-cm dishes, and subjected to three different drug selections: G418 only, hygromycin only, and G418 plus hygromycin. On Day 9, ESC colonies were picked and divided into two groups: one for lysate preparation for PCR analysis of the deletion events and the other for cell culture to make frozen stocks.

### RNA-seq

The total RNA was extracted with RNeasy Plus Mini Kit (Qiagen). MEF feeder cells were removed from ESC culture before RNA extraction by plating cells on a gelatin-coated dish for 30 min during the passaging and expanding unattached cells in a new dish. Library preparation was performed using the TruSeq stranded mRNA sample prep kit (Illumina) according to the manufacturer’s instructions. Sequencing was performed on the Illumina NovaSeq 6000 platform in a 100 bp paired-end mode. Sequenced reads were mapped to the mouse reference genome sequences (mm10) using TopHat v2.0.13 (Trapnell *et al*, 2009) in combination with Bowtie2 ver. 2.2.3 (Langmead & Salzberg, 2012) and SAMtools ver. 0.1.19 (Li *et al*, 2009). The fragments per kilobase of exon per million mapped fragments (FPKMs) was calculated using Cufflinks version 2.2.1 (Trapnell *et al*, 2010) (Supplementary Table 1).

### Data availability

The RNA-seq data are available in the DNA Data Bank of Japan (DDBJ) Sequencing Read Archive under the accession numbers DRA013360.

## AUTHOR CONTRIBUTIONS

Conceptualization, K.H.; Methodology, K.H; Investigation, M.M., J.Y., I.T., K.H.; Writing the original draft, K.H., M.M.; Resources, K.H.; Supervision, K.H.; Project administration, K.H.; Funding acquisition, K.H.

## ACKNOWLEDGMENTS

We acknowledge the NGS core facility of the Genome Information Research Center at the Research Institute for Microbial Diseases of Osaka University for support with RNA sequencing and data analysis. We thank Dr. Kosuke Yusa for providing the plasmids pPGKneo-F2F and pPGKhyg-F2F. This work was supported by Grants-in-Aid for Scientific Research from the Ministry of Education, Culture, Sports, Science, and Technology of Japan (JP16H04683, JP18K19275, JP20H03174 for K.H.). This work was also supported in part by Nara Medical University Grant-in-Aid for Collaborative Research Projects and a research grant from the Takeda Science Foundation (K.H.), Naito Foundation (K.H.), and Daiichi Sankyo Foundation of Life Science (K.H.).

